# A Yeast Surface Display Platform for Screening Dimeric Mammalian Receptors

**DOI:** 10.64898/2026.01.29.702702

**Authors:** Ethan W. Slaton, Elise C. Krivanek, Blaise R. Kimmel

## Abstract

Discovering proteins that modulate receptor activity remains a key challenge in the field of protein design and engineering. Traditionally, identifying proteins that interact with receptors often relies on binding as a selection criterion, yielding limited information about the function of discovered binders in a library, including the ability to activate or block signaling cascades associated with the receptor of interest. As a result, extensive downstream characterization is required to assess the biological relevance of discovered binders. To address this issue, we have developed a high-throughput screening system to screen dimeric mammalian receptors using yeast surface display. We demonstrate the programmed dimerization of the extracellular domains of mammalian receptors in yeast via engineered induction pathways, thereby enabling receptor expression and the secretion of associated native cytokines. This surface expression of the involved subunits for the protein receptor and cytokine-induced dimerization activity indicates that the receptor has been activated and is expected to trigger a DNA-driven signaling cascade within a mammalian cell. This system provides a modular platform technology that advances existing yeast-display systems, demonstrating the effectiveness of these high-throughput platforms for screening the function of mammalian receptors. This work is expected to provide a rapid, cost-effective approach to the molecular discovery of novel biologics for targeting dimeric mammalian receptors.

## Introduction

Receptors are involved in several biological processes, including immune responses, cell-to-cell signaling, and cellular proliferation, making these proteins important therapeutic targets.^1–3^ While cell-surface receptors are key targets for immunotherapies, identifying protein-based modulators that alter receptor function in a high-throughput manner remains a significant challenge.^4,5^ This challenge is exacerbated by the limitations of developing and working with high-throughput mammalian-based systems, which require aseptic cellular conditions for materials used in mammalian cell culture, and by the high costs associated with cell maintenance, necessitating several months of effort to generate an evolved protein binder for iterative protein evolution campaigns.^6,7^ Researchers prefer to use mammalian cells over other cell-based models as these biological systems more accurately bridge the gap between *in vitro* and *in vivo* experimentation, particularly due to mammalian cells maintaining accurate biological signaling pathways associated with cell-receptor disturbances (e.g., binding events), as well as proper protein expression and post-translational modification (PTM) events (e.g., glycosylation).^8,9^ One potential strategy to address the challenges of mammalian high-throughput screens is to use yeast. Yeast, and specifically yeast surface display, have been used to create platforms that enable high-throughput screening of large libraries against targets such as proteases and G protein-coupled receptors (GPCRs).^10–12^ Furthermore, yeast and mammalian cells share substantial similarities in metabolic processes and signaling pathways, including the mitogen-activated protein kinase (MAPK) pathway.^13,14^ While effective at screening large libraries and modeling some pathways exhibited in mammalian cells, yeast has a different glycosylation pattern when compared to mammalian cells, which is often essential for functional recombinant proteins.^15,16^ This alteration in the glycosylation pattern could lead to translation issues when testing yeast-discovered therapeutics in mammalian cells. Despite this limitation, the advantages of using yeast for high-throughput screenings motivate the development of novel phenotypic screens to discover potential therapeutics.

To overcome the challenges associated with mammalian cell culture and the expression of functional protein receptors in yeast, we have developed a high-throughput yeast surface display system for screening large protein libraries to identify receptor modulators. This system uses endoplasmic reticulum (ER) sequestration to facilitate interactions between receptor components and modulators within the ER.^17^ After this interaction, the receptor is trafficked to the yeast surface and is covalently attached via disulfide bonds formed between Aga1 and Aga2. Once on the yeast surface, high-affinity recognition sequences (e.g., c-Myc, FLAG, HA) are adjacent to the receptor motif, allowing direct visualization of receptor display efficiency before and after ligand binding and providing a quantitative strategy to identify the presence of receptor components **(Figure 1)**. Once displayed, the system provides a high-throughput approach for distinguishing ligand binding from receptor dimerization, enabling the isolation of both agonistic and antagonistic ligands for receptors of interest. We hypothesize that this dimerization event, induced by the yeast modulator, will be sufficient to generate a signal cascade when the modulator is purified and introduced into mammalian cell culture. In this work, we describe the development, optimization, and evaluation of the system for a model receptor-cytokine pair, focusing on IL-7 and IL-21, to develop and screen potential therapeutic binders for IL-7Rα and IL-21R.

**Figure 1:**
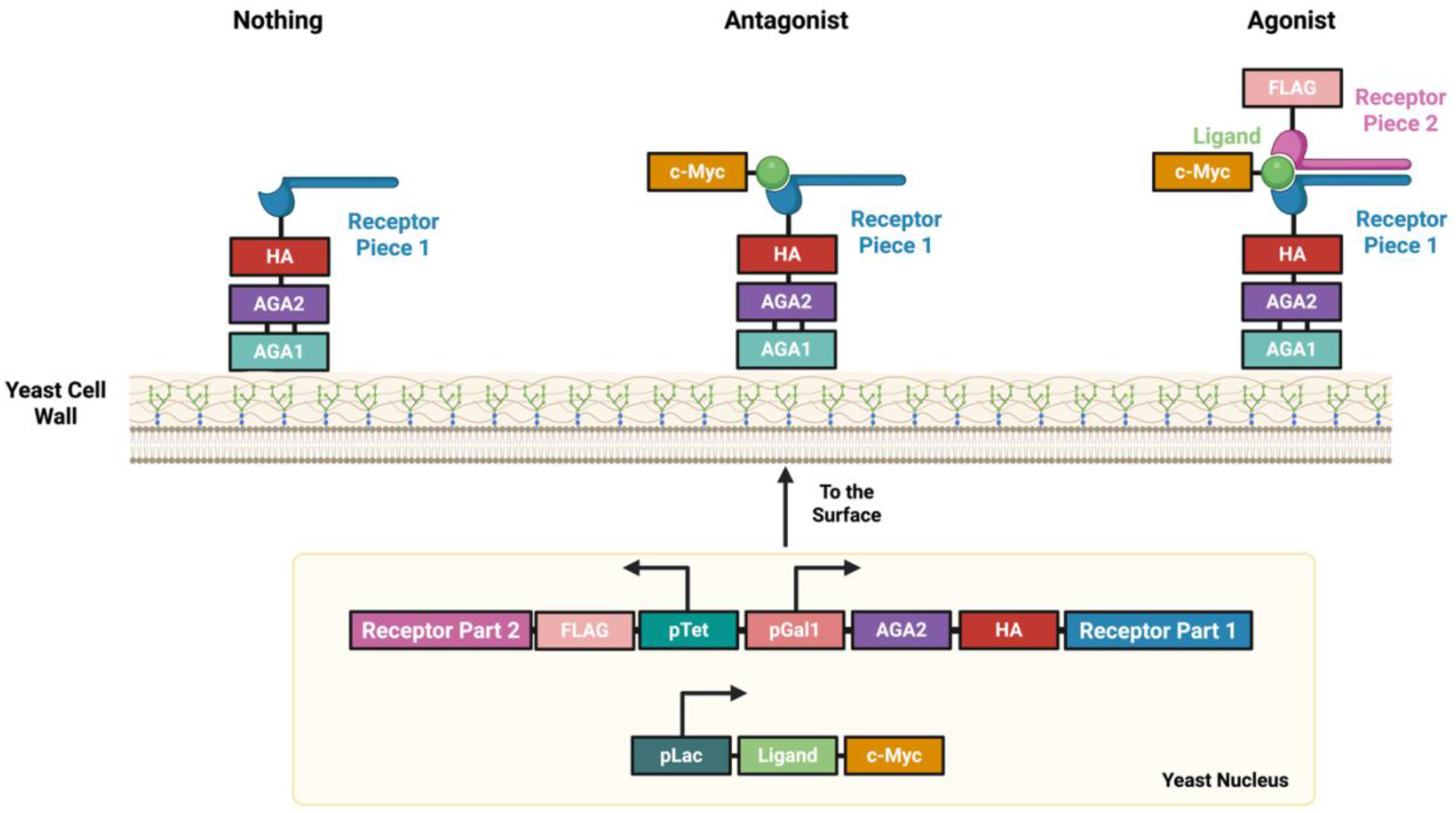
A high-throughput yeast surface display system to identify antagonists and agonists from large libraries. We observe three distinct outcomes on this screen after receptor parts 1 and 2 interact with the ligand in the yeast ER. These results depend on the ligand-bound effects on receptor dimerization. If the c-Myc tag is not present on the surface, as this indicates that the ligand in that cell did not bind to the receptor well, if the c-Myc tag is present, but the FLAG tag is not detectable, suggesting that this ligand could potentially be an antagonist, and if the c-Myc and FLAG signals are present, meaning that the ligand could act as an agonist.

## Results and Discussion

### Surface Display of IL-7Rα

To begin this work, we constructed pEWS1 **(Figure 2A)**, a plasmid derived from the previously constructed pY2 plasmid (Martinusen *et al*.).^18^ pEWS1 replaces the *β-Estradiol* promoter within pY2 with a tetracycline (Tet) promoter (pTet7).^19^ This replacement enabled us to create a bidirectional plasmid, in which one side is inducible under the pTet7 promoter and the other is inducible under a galactose promoter (pGal1). The pGal1 side of this plasmid contains Aga2, which will serve as the anchor to the yeast surface. Importantly, each side of the plasmid contains unique Golden Gate assembly cut sites, BsmBI and BsaI, providing a modular platform for testing interactions between different receptor pieces. To test the ability of *S. cerevisiae* to express the extracellular portion of mammalian receptors correctly, the extracellular portion of mouse IL-7Rα, which serves as one piece of the dimer that forms IL-7R, was purchased. IL-7Rα was inserted into the pGal1 side of the pEWS1 plasmid via a BsmBI Golden Gate assembly **(Figure 2B)**. IL-7Rα was flanked with two epitope tags (i.e., FLAG, HA), which can be stained with fluorescent antibodies when the construct is anchored to the surface, to validate both the expression of the protein and characterize adequate display on the yeast cell wall, ensuring that IL-7Rα is being folded properly within the yeast cells. If IL-7Rα were to be misfolded, the epitope tag on the C-terminal of the construct would likely also be misfolded, leading to a loss in fluorescent signal after staining when running flow cytometry.

**Figure 2:**
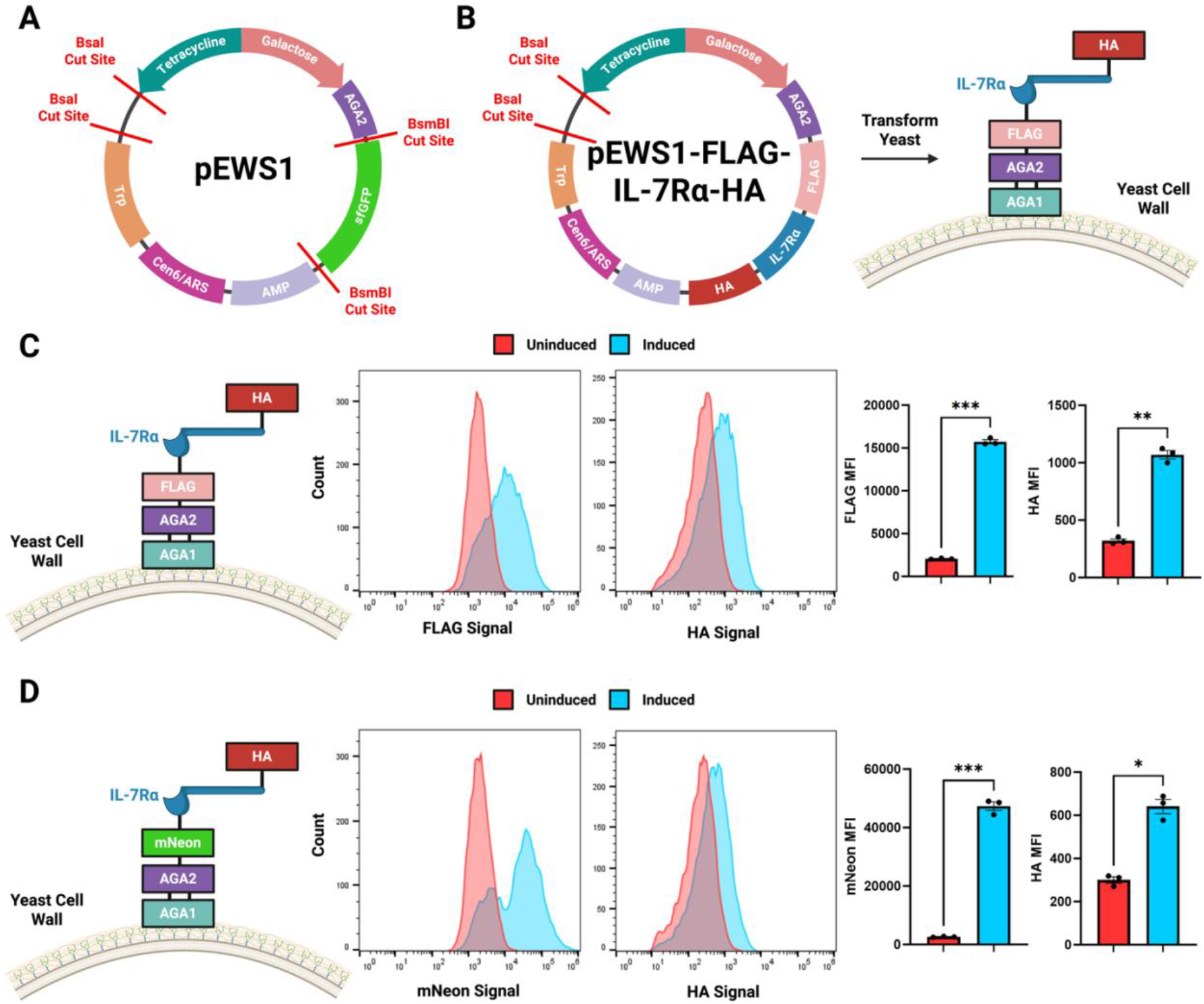
Surface Displaying IL-7Rα. **A)** The pEWS1 plasmid used to display mammalian receptor pieces on the yeast surface. **B)** The pEWS1-FLAG-IL-7Rα-HA that was constructed following a BsmBI Golden Gate, allowing for the expression of IL-7Rα on the yeast surface. **C)** The protein construct expressed on the yeast surface following galactose induction of pEWS1-FLAG-IL-7Rα-HA, with flow cytometry data comparing the uninduced and induced populations of this construct are included (n=3). **D)** The protein construct expressed on the yeast surface when the FLAG epitope tag is replaced with an mNeon fluorescent protein. Flow cytometry data comparing the uninduced and induced populations of this construct are included (n=3). *P* values were determined by using a paired t-test between the uninduced and induced samples (**C, D**). All data are shown as mean±s.e.m.

After creating the pEWS1-FLAG-IL-7Rα-HA construct and confirming successful cloning into the plasmid, the engineered vector was transformed into the triple-transcription factor system, referred to as BIT (containing the *β-Estradiol, IPTG, and tetracycline* transcription factors). It should be noted that the *B-Estradiol* transcription factor was not used in this work. This strain is crucial for creating a tri-orthogonal system in yeast, enabling user-directed control of the plasmid for modular integration and expression of three-protein components, which are typically necessary for protein-receptor dimerization. As shown in the histograms and supported by mean fluorescence intensity (MFI) bar graphs **(Figure 2C)**, a notable increase in both the FLAG and HA signals was observed when comparing the uninduced and induced populations. This indicates that the extracellular portion of IL-7Rα is folded correctly and expressed on the yeast surface.

Once this was successful, we proceeded to express the extracellular IL-7Rα with an mNeonGreen (mNeon) fluorescent protein **(Figure 2D)**. This choice was made to use a split mNeon fluorescent protein in RADAR. The split mNeon emits fluorescence when the two independent fragments reconstitute into a functional fluorophore.^20^ We hypothesized that the expression of a split fluorescent protein could provide a robust strategy for determining whether IL-7Rα dimerizes with IL-2Rγ to form the complete IL-7R complex, as an alternative to traditional epitope tags. A potential challenge in using a fluorescent protein, rather than using epitope tags on individualized receptor pieces, was how to design the system for effective surface display of the fusion protein complexes (e.g., split fluorescent proteins with IL-7R subunits), given the bulkiness of the fluorescent protein compared to an epitope tag (∼10x greater molecular weight for the fluorescent protein compared to the epitope tag). To investigate this, we replaced the FLAG epitope tag with the complete mNeon protein, forming the pEWS1-mNeon-IL-7Rα-HA construct. When assembled, this plasmid was transformed into the BIT yeast strain and expressed effectively. From the flow cytometry data in **Figure 2D**, we observed significant mNeon and HA signals (*P*=0.001 and *P*=0.0179, respectively) when comparing the uninduced and induced populations, suggesting that the mNeon-IL-7Rα-HA construct was successfully displayed on the yeast surface.

### Testing Dimerization of IL-7Rα and IL-2Rγ with IL-7

After validating successful surface display of IL-7Rα on yeast, we evaluated whether IL-7 could induce dimerization between IL-7Rα and IL-2Rγ, thereby mimicking the initiating factors in the IL-7R activation pathway in mammalian cells **(Figure 3)**. To do this, we ordered three different gene fragments: HA-IL-7Rα-mNeon-11, FLAG-IL-2Rγ-mNeon-1-10, and IL-7-cMyc. The HA-IL-7Rα-mNeon-11 and FLAG-IL-2Rγ-mNeon-1-10 fragments were inserted into the pEWS1 plasmid **(Figure 3A)**, and the IL-7-cMyc fragment was inserted into a receiver plasmid under the pLacFec promoter (for IPTG induction) **(Figure 3B)**. The rationale for using split mNeon was to avoid the need for fluorescent antibodies to stain surface epitope tags and to provide a direct method for determining whether both receptor pieces are present on the surface. As a safeguard, the HA tag was included on the IL-7Rα fragment to validate the yeast surface display of the protein receptor. Similarly, the FLAG tag was included in the IL-2Rγ fragment to enhance the quantification and visualization of the attachment of the IL-2Rγ fragment to the IL-7Rα portion of the receptor, in case the split mNeon did not reconstitute to produce a signal. Additionally, we placed a c-Myc tag on the IL-7 cytokine to simultaneously determine ligand binding to the receptor upon receptor dimerization on the yeast surface.

**Figure 3:**
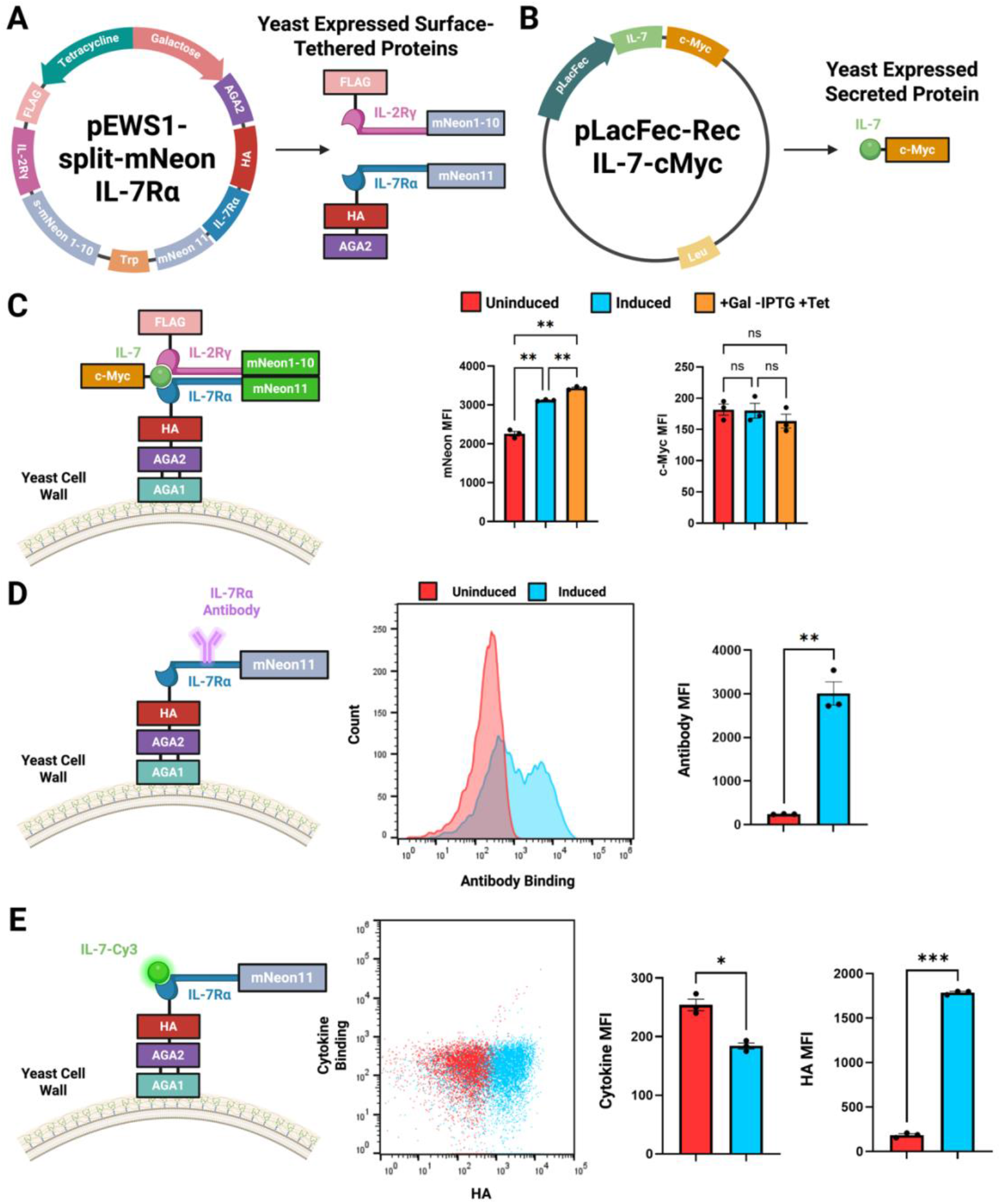
IL-7Rα Expression and Testing on the Yeast Surface. **A)** The pEWS1-split-mNeon-IL-7Rα plasmid was created from pEWS1 to express the IL-7Rα and IL-2Rγ that create the IL-7R receptor upon dimerization. **B)** The pLacFec (IPTG inducible) receiver plasmid used to express IL-7. **C)** The protein construct expressed on the yeast surface following expression of IL-7Rα, IL-2Rγ, and IL-7, if the receptor dimerizes in the presence of IL-7 containing the split mNeon protein. Flow cytometry data comparing the cell populations when uninduced, fully induced, and in the absence of IL-7 is shown (n=3). **D)** Expression of IL-7Rα on the yeast surface followed by anti-IL-7Rα staining. Flow cytometry data comparing antibody binding when IL-7Rα is expressed and not expressed on the yeast surface is shown (n=3). **E)** Expression of IL-7Rα on the yeast surface followed by IL-7-Cy3 binding, where flow cytometry data comparing IL-7 binding when IL-7Rα is expressed and not expressed on the yeast surface is shown (n=3). *P* values were determined by using a repeated measure one-way ANOVA (**C**) or using a paired t-test between the uninduced and induced samples (**D, E**). All data are shown as mean±s.e.m.

Following the cloning of these fragments and subsequent sequencing, these plasmids were transformed into the BIT yeast strain for induction. We experimentally validated the triple-orthogonal system (galactose, IPTG, and tetracycline) and observed that the mNeon fluorescent signal was highest, regardless of IL-7 ligand expression, relative to uninduced populations lacking either receptor motif **(Figure 3C)**. This ligand-independent dimerization of the receptor system is expected due to the high binding affinity and recognition between the two subunits of the split mNeon protein, which were expressed at high concentrations in the ER in our yeast system. Furthermore, these results indicate that the split mNeon fragments are functionally capable of binding and producing a measurable signal, validating high expression of both the IL-7Rα and IL-2Rγ subunits within the cells. While we expect to express adequate levels of the IL-7 ligand within the yeast cell system, we were unable to detect significant levels of the c-Myc-tagged IL-7 cytokine on the surface, potentially indicating low IL-7 binding to the surface-displayed IL-7R. Due to the ligand-independent dimerization of the split mNeon, further testing of mammalian receptor dimerization will be conducted without the use of the split mNeon fragment.

### Yeast Glycosylation Altering the Binding Affinity of IL-7

We next examined the ability of yeast-expressed IL-7 to bind the IL-7R complex, initially observing limited fluorescent signal from the c-Myc tag linked to the IL-7 ligand. To evaluate this, we purchased a fluorescent antibody that binds to IL-7Rα and a commercially available IL-7 cytokine to assess whether the IL-7Rα receptor expressed on the yeast surface was folded equivalently to that expressed in mammalian cells.^21,22^ For testing the ability of the IL-7Rα antibody to bind, we induced the expression of the IL-7Rα portion of the pEWS1-split-mNeon-IL-7R plasmid using galactose to attach the receptor to the surface **(Figure 3D)**. The yeast cells were then stained with the IL-7Rα antibody, and we observed strong antibody binding, indicating that the IL-7Rα portion of the receptor is folding correctly and exhibits structural similarity to a mammalian receptor. We note that this result is promising, as it demonstrates that yeast can correctly fold the extracellular portion of IL-7Rα.

Following this, we stained the IL-7 cytokine with a Cy3-NHS ester dye, a commercially available reagent traditionally used to label surface-exposed lysine residues on proteins. Following the experimental protocols for IL-7Rα surface display, the IL-7Rα piece of the pEWS1-split-mNeon-IL-7R plasmid was expressed on the yeast surface. The IL-7-Cy3 cytokine was then incubated with the cells to bind to IL-7Rα **(Figure 3E)**. The HA tag was also stained with a fluorescent antibody in this experiment to confirm that IL-7Rα was expressed on the cell surface. From our flow cytometry data, we observe minimal binding of the IL-7 cytokine to IL-7Rα on the yeast surface. Therefore, we hypothesized that this was due to IL-7 being highly dependent on proper glycosylation of the IL-7R subunits, and that the glycosylation pattern would interfere with binding affinity for IL-7 ligands. As noted by McElroy *et al*., the K_d_ value of IL-7 to glycosylated IL-7Rα (CHO) was 62 nM, while unglycosylated IL-7Rα (*E. coli*) was just 18 µM – confirming our prediction.^23^ Since glycosylation patterns differ between yeast and mammalian cells, we expect this post-translational modification to be a key requirement for the development of cell-specific tolerated ligands. For our technology, this represents a potential limitation for the use of mammalian-derivatized ligands that bind to mammalian- and yeast-produced receptors. This result positions RADAR and yeast, in general, as particularly suited to identifying potential agonists and antagonists of mammalian receptors that are less dependent on glycosylation, motivating our subsequent experiments with IL-21R.

### Testing Dimerization of IL-21R and IL-2Rγ with IL-21

After completing preliminary tests using mouse IL-7Rα in our system, we shifted our focus to an interleukin that is less dependent on glycosylation: mouse IL-21R. In a previous study, this receptor was expressed in *E*.*coli* and following purification, successfully formed a complex with IL-21, suggesting it is less reliant on glycosylation.^24^ Similar to the IL-7R dimerization experiment in **Figure 3**, we ordered gene fragments for IL-21R and IL-21 and inserted them into pEWS1 **(Figure 4A)** and pLacFec-Rec **(Figure 4B)**, respectively. Following cloning and sequencing, these plasmids were co-transformed into the BIT yeast strain for induction **(Figure 4C)**. The results of this experiment show that IL-2Rγ and IL-21R dimerize in the absence of IL-21 induction. This result mirrored the results obtained with IL-7R using the split mNeon, which was unexpected. To confirm folding of IL-21R, we purchased an antibody that binds to IL-21R and induced only the IL-21R portion of the pEWS1-IL21R plasmid **(Figure 4D)**. Following induction, the yeast cells were stained with the IL-21R antibody. We observed a significant shift in antibody fluorescence in the induced population compared with the uninduced population. This shift was smaller than expected, but due to the antibody binding near or around the IL-21 binding site within IL-21R, we hypothesize that glycosylation differences between yeast and mammalian receptors could hinder antibody binding. We also purchased a commercial IL-21 cytokine and stained it with a Cy3-NHS ester dye to test IL-21 binding to yeast-expressed IL-21R. IL-21R was induced from the pEWS1-IL-21R plasmid, and IL-21-Cy3 was then incubated with cells to test binding **(Figure 4E)**. From this result, we observe a significant increase in Cy3 fluorescence in the induced population, indicating that IL-21 can bind to yeast-expressed IL-21R.

**Figure 4:**
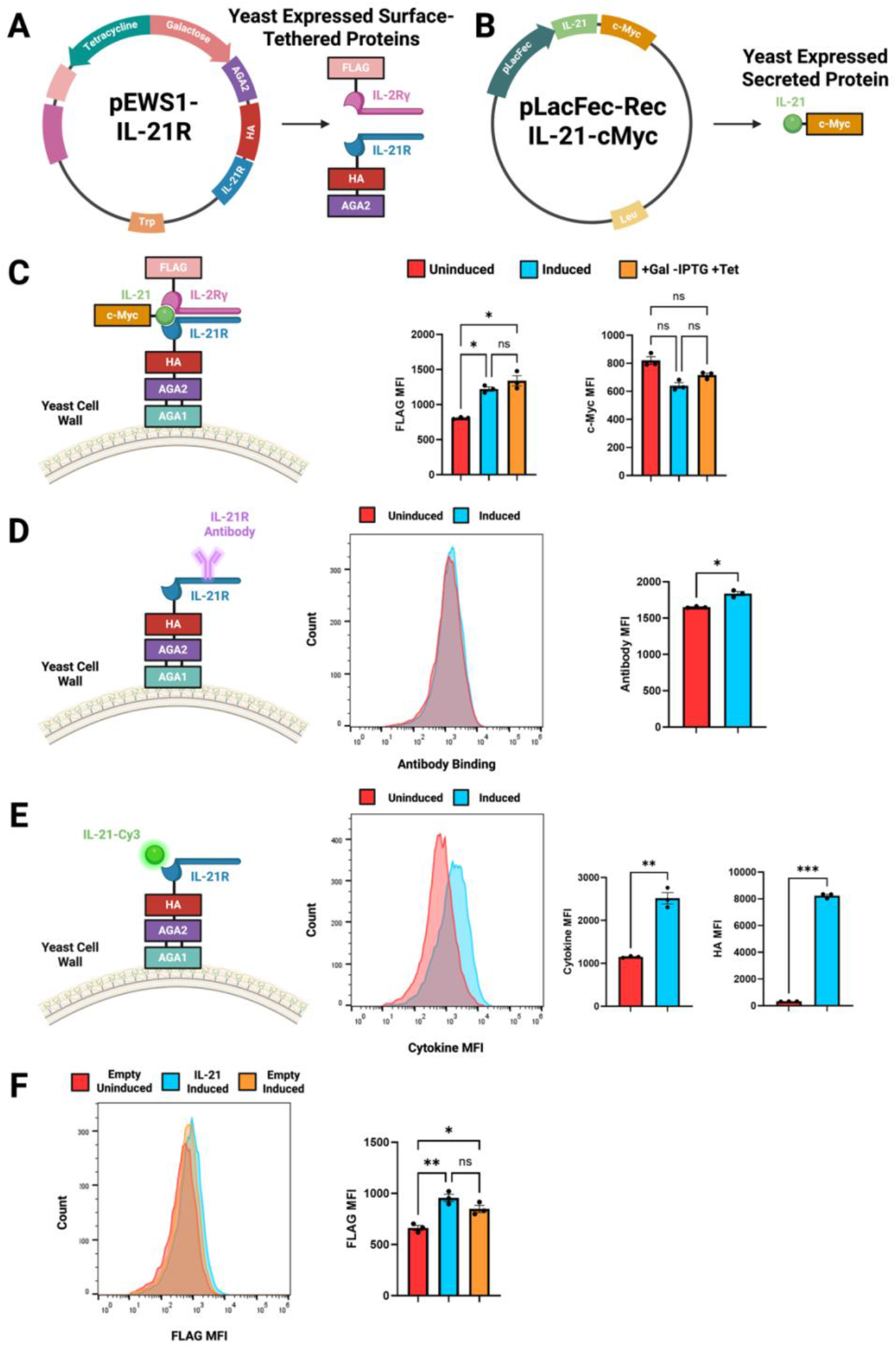
IL-21R Expression and Testing on the Yeast Surface. **A)** The pEWS1-IL-21R plasmid created from pEWS1 to express the IL-21 and IL-2Rγ that create the IL-21 receptor upon dimerization. **B)** The pLacFec (IPTG inducible) receiver plasmid used to express IL-21. **C)** The protein construct expressed on the yeast surface following expression of IL-21R, IL-2Rγ, and IL-21, with flow cytometry data comparing the cell populations when uninduced, fully induced, and in the absence of IL-21 is shown (n=3). **D)** Expression of IL-21R on the yeast surface followed by anti-IL-21R staining, with flow cytometry data comparing antibody binding when IL-21R is expressed and not expressed on the yeast surface is shown (n=3). **E)** Expression of IL-21R on the yeast surface followed by IL-21-Cy3 binding, with flow cytometry data comparing IL-21 binding when IL-21R is expressed and not expressed on the yeast surface is shown (n=3). **F)** The pEWS1-IL-21R plasmid was transformed into the BIT yeast strain with either an empty pLacFec-Rec or the pLacFec-IL-21-cMyc plasmid, with flow cytometry data comparing the uninduced empty (red), induced IL-21 (blue), and induced empty (orange) are included (n=3). *P* values were determined using an ordinary one-way ANOVA (**C, F**), or by paired t-test between the uninduced and induced samples (**D, E**). All data are shown as mean±s.e.m.

Based on our understanding that IL-21 can bind to IL-21R expressed in yeast, we wanted to explore why the receptors dimerized even in the absence of IL-21 induction in **Figure 4C**. We hypothesized that this unexpected dimerization could result from the pLacFec promoter being leaky and expressing small amounts of IL-21 in the absence of induction. To test this, we co-transformed the pEWS1-IL-21R plasmid with an empty pLacFec-Rec plasmid containing no IL-21 and compared that to cells containing pEWS-IL-21R and pLacFec-Rec-IL-21-cMyc **(Figure 4F)**. This result shows that the IL-21R and IL-2Rγ appear to dimerize within the yeast ER without the need for IL-21. We hypothesize that this dimerization is due to the high concentration of both receptor components within the yeast ER, and incorporating a weaker promoter in place of the pTet7 may resolve this problem. Additionally, we believe that the difference in MFI between the empty and IL-21 samples, while not significant, could be enough to enrich the IL-21 population in a mock sort with a mixed population of empty and IL-21 samples.

## Conclusions

We present a high-throughput functional screen to identify novel binders to dimeric mammalian receptors in yeast. These data from our work suggest that yeast can effectively display mammalian receptors on its surface and bind interleukins, depending on the receptor used. The antibody and cytokine binding experiments with IL-7Rα and IL-21R suggest that yeast can fold the mammalian receptor subunits properly; however, the glycosylation patterns on these receptors will differ from those in mammalian cells, and some may form complexes with the native cytokines in yeast^15^. This could severely limit the ability of this screen to isolate modulators that depend on mammalian glycosylation patterns to bind. On the other hand, this application will be particularly valuable for screening cancer therapeutics to expand the functionality of proteins from being receptor binders for drug and gene delivery (e.g., nanobody therapies)^5,25,26^ and the impact these delivery agents may play in enhancing therapeutic cargo delivery in cancer nanotechnology (e.g., functionalized proteins on the surface of lipid nanoparticles^27^ and polymersomes^28^), as well as extending the role that these binders play in altering cellular functions and signaling as agonistic and antagonistic therapies^29^. This is due to most cancer cells having high mutation rates, which can alter glycosylation patterns on receptors expressed on those cells. If glycosylation-independent protein therapeutics were to be discovered as selective binders against these receptors, and where the proteins could functionally alter the signaling cascade of the bound receptor motif, these functional protein-based therapies may have the potential to overcome the high mutation rates in cancer cells, allowing the therapeutics to be widely effective in the tumor microenvironment.

Moving forward, we plan to conduct a mock sort between IL-21 and empty-receiver populations to determine whether a weaker promoter is needed to utilize the system^10^. In parallel, we will continue to optimize the system by using weaker promoters to better control the concentration of receptor fragments within the yeast ER, thereby minimizing what appears to be cytokine-independent dimerization. Following this, we will screen large nanobody libraries to identify novel agonists and antagonists targeting IL-7Rα and IL-21R. Subsequently, we will test these nanobodies in IL-7R- and IL-21R-reporter strains to verify the functionality of the receptors used in the yeast-based screen on natively expressed receptor variants in mammalian cells. If successful, we will proceed to test the isolated nanobodies in mouse models for potential therapeutic applications. The impact of this advancement will shape throughput for screening functional protein binders, enabling the expression of mammalian receptor variants in yeast-based systems to accelerate the discovery of novel protein therapeutics and minimize the costs of mammalian cell culture in protein discovery^30^.

## Materials and Methods

### pEWS1 Plasmid Construction

The pY2 plasmid was a gift from Dr. Carl Denard (Addgene#218115). One side of this bidirectional plasmid contains Aga2 under a pGal1 promoter, which was used as the anchor for surface expression. The other side of this plasmid contains a 3PminCYC1(lexA-box) promoter. To construct pEWS1, the entire pY2 plasmid was PCR’d, besides the lexA-box and its complementary signal peptide. This linear DNA fragment was then combined with a pTet7 promoter that was PCR-amplified to include complementary overhangs for HiFi assembly (NEB #E2621S). The pTet7 promoter was a gift from Dr. Tom Ellis (Addgene#180668). Once this plasmid was made, almost the entire plasmid was PCR’d again, and a gene fragment containing the signal peptide for surface display was inserted via HiFi assembly following the pTet7 promoter.

### pLacFec-Rec Plasmid Construction

The pLacFec-Rec plasmid was created using pieces from the MoClo Yeast Toolkit (Addgene #1000000061) as well as pieces from the Multiplex Yeast Toolkit (Addgene #1000000229). The kits were gifts from Dr. John Dueber and Dr. Tom Ellis, respectively. These kits are complementary and work by selecting kit components to create plasmids using BsaI (NEB #R3733L) Golden Gate technology. The only component used in the construction of this plasmid that was not already part of either kit was the receiver insert. This receiver insert contains BbsI (NEB #R3539L) Golden Gate sites for inserting DNA of interest into the receiver. This piece was ordered as a gene fragment and contained BsaI overhangs, which allowed it to be considered a piece three in these kits. To construct the plasmid, a BsaI Golden Gate reaction was conducted containing pYTK002, pMYT005, the receiver inserts, pYTK056, pYTK072, pYTK075, pYTK081, and pYTK083. Following the Golden Gate reaction, the plasmid was transformed into NEB 10-beta competent E. coli (NEB #C3019H) and plated on ampicillin selection plates (100 µg/mL) for approximately 12-18 hours.

### Cloning into pEWS1

Once the pEWS1 plasmid was constructed, gene fragments were ordered to insert receptor pieces containing BsaI (NEB #R3733L) golden gate cut sites and the respective overhangs (for insertion into the pTet7 side of the plasmid) or BsmbI (NEB #R0739L) cut sites and the respective overhangs (for insertion into the pGal1 side of the plasmid). Following the Golden Gate process, the plasmids were transformed into NEB Turbo Competent E. coli (NEB #C2984H), which were made competent using the Zymo Research Mix & Go! *E. coli* Transformation kit (Zymo #T3001) and plated on ampicillin selection plates (100 µg/mL) for approximately 12-18 hours. Colonies were then miniprepped using the Qiagen QIAprep Spin Miniprep Kit (Qiagen #27104). The sequence of the constructs was then confirmed using Genewiz Plasmid-EZ whole plasmid sequencing.

### Cloning into pLacFec-Rec

Once the pLacFec-Rec plasmid was constructed, gene fragments were ordered to insert DNA containing BbsI (NEB #R3539L) golden gate cut sites and the respective overhangs. Following the Golden Gate process, the plasmids were transformed into NEB Turbo Competent E. coli (NEB #C2984H), which were made competent using the Zymo Research Mix & Go! *E. coli* Transformation kit (Zymo #T3001) and plated on ampicillin selection plates (100 µg/mL) for approximately 12-18 hours. Colonies were then miniprepped using the Qiagen QIAprep Spin Miniprep Kit (Qiagen #27104). The sequence of the constructs was then confirmed using Genewiz Plasmid-EZ whole plasmid sequencing.

### BIT Yeast Strain Creation

To create the BIT yeast strain, plasmids were constructed using the MoClo Yeast Toolkit (Addgene #1000000061) as well as pieces from the Multiplex Yeast Toolkit (Addgene #1000000229) that contained a constitutive promoter, the respective transcription factor, and a terminator. The transcription factors enable the use of the β-Estradiol, IPTG, and Tetracycline promoters included in the kit. Once three plasmids were constructed, one for each transcription factor, the promoter, transcription factor, and terminator were PCR’d with unique BsaI Golden Gate cut sites. The three PCRs were then inserted into the MET-HYGRO integration vector (Addgene#218113), a gift from Dr. Carl Denard. The vector was then linearized using NotI cut sites and inserted in the yeast chromosome via lithium acetate cell transformation as discussed by Martinusen *et al*.^18^

### Yeast Transformations

The BIT yeast strain was previously constructed from the EBY100 yeast strain. The BIT strain contains the transcription factors necessary to utilize the IPTG, Tetracycline, and β-Estradiol promoters within the Multiplex Yeast Toolkit (Addgene #1000000229). Competent BIT cells were then created using the Zymo Research Frozen-EZ Yeast Transformation II Kit (Zymo #T2001). Following sequence verification of plasmids, they were transformed into the competent BIT cells and plated on selection plates (YNB CAA 2% GLU or SC-Leu-Trp 2% GLU) for 2-4 days (30°C, 250 rpm).

### Cytokine Staining

The ILs were purchased from Biolegend (#577802/#574502). The ILs were reacted for 4-6 hours with a Cy3-NHS ester dye (AAT Bioquest #141) in 15-fold excess. Following the reaction, dialysis was performed overnight in PBS (pH 7.4). After dialysis, the dyed IL was concentrated in Amicon Ultra Centrifugal Filters (Sigma #2025-08-01) with several washes until all excess dye was removed.

### Yeast Induction for Protein Expression

Colonies that grew on selection plates were inoculated in 2mL of selection media (YNB CAA 2% GLU or SC-Leu-Trp (Sigma #Y0750) 2% GLU) for 18-24 hours at 30°C (250 rpm) to reach saturation. Optical densities (OD_600_) were taken when the cultures reached saturation, and new cultures were inoculated at an OD of 0.5. These cultures were outgrown (30°C, 250 rpm) until they reached an OD_600_ between 2-4. Once this OD_600_ range was reached, the culture volume required to reach an OD_600_ of 0.5 was calculated. The media that was used to induce contained some combination of YNB CAA 2% GAL, SC-Leu-Trp 2%-GAL, 50 mM IPTG (Fisher #BP1755-100), and 100 ng/mL anhydrotetracycline hydrochloride (Sigma #37919). Uninduced samples were inoculated in either YNB CAA 2% GLU or SC-Leu-Trp 2% GLU. Samples for induction were grown for 12-16 hours at 30°C at 250 rpm.

### Flow Cytometry

Following induction, the OD_600_ was measured, and the amount of culture required to obtain 2 million yeast cells was calculated. This amount of culture was washed in PBS 0.5% BSA (Goldbio #9048-46-8). The cells were then stained at room temperature for 90 minutes and stained with the appropriate antibodies or cytokine: anti-HA Alexa 594 (Biolegend #901511), anti-c-Myc Alexa 555 (Thermo Fisher #MA1-980-A555), anti-FLAG Alexa 488 (Biolegend #637317) for the IL-7 experiments, or anti-FLAG PerCP/Cyanine5.5 (Biolegend #637325) for IL-21 experiments. IL-21-Cy3 staining was conducted at 100 nM, and IL-7-Cy3 staining was conducted at 120 nM. PE anti-IL-21R (Biolegend #131905) and PE anti-IL-7Rα (Biolegend #135009) were used for staining receptor pieces. Following the 90 min stain, cells were washed and resuspended in PBS 0.5% BSA for flow cytometry (Attune NxT Acoustic Focusing Cytometer). For flow cytometry data analysis FlowJo was utilized. Statistical analysis and plot creation were conducted using GraphPad Prism.

## AUTHOR INFORMATION

### Author Contributions

Ethan Slaton: Wrote the original draft of the manuscript, generated all figures and graphics for the manuscript, edited, revised, and approved the final version of the manuscript. Elise Krivanek: Supported experimental design and data generation for the manuscript. Blaise Kimmel: Wrote the original draft of the manuscript, edited, revised, and approved the final version of the manuscript, and acquired funding to support the work.

## Acknowledgements

This work was supported in part by The Ohio State University Center for Cancer Engineering-Curing Cancer Through Research in Engineering and Sciences. B.R.K. acknowledges financial support from the Prostate Cancer Foundation Young Investigator Award. We thank Natalie Clay for insightful discussions and support in the revisions of this work. We thank the laboratory of Dr. Carl Denard for sharing the pY2 plasmid (Addgene #218115) and the MET-HYGRO integration vector (Addgene #218113). All figures created with Biorender.com.

