## Supplemental Information for "A Yeast Surface Display Platform for Screening Dimeric Mammalian Receptors"

### Supplemental Figures

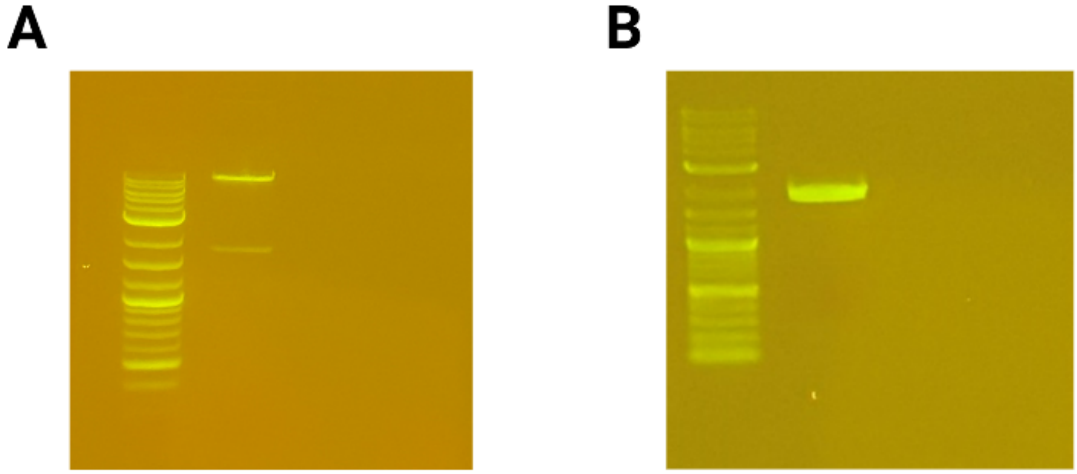


**Supplemental Figure 1: BIT Transcription Factor Integration. A)** A DNA gel showing the NotI digestion of the MET-Hygro plasmid containing the BIT constitutive promoters, transcription factors, and terminators. The top band was gel purified and integrated into the yeast chromosome. **B)** Following integration, genomic DNA from a colony that grew on selection media was obtained. A PCR was then run to confirm integration with one primer annealing to the genomic locus outside of the homology arm and one primer annealing to a portion of the integrated construct.

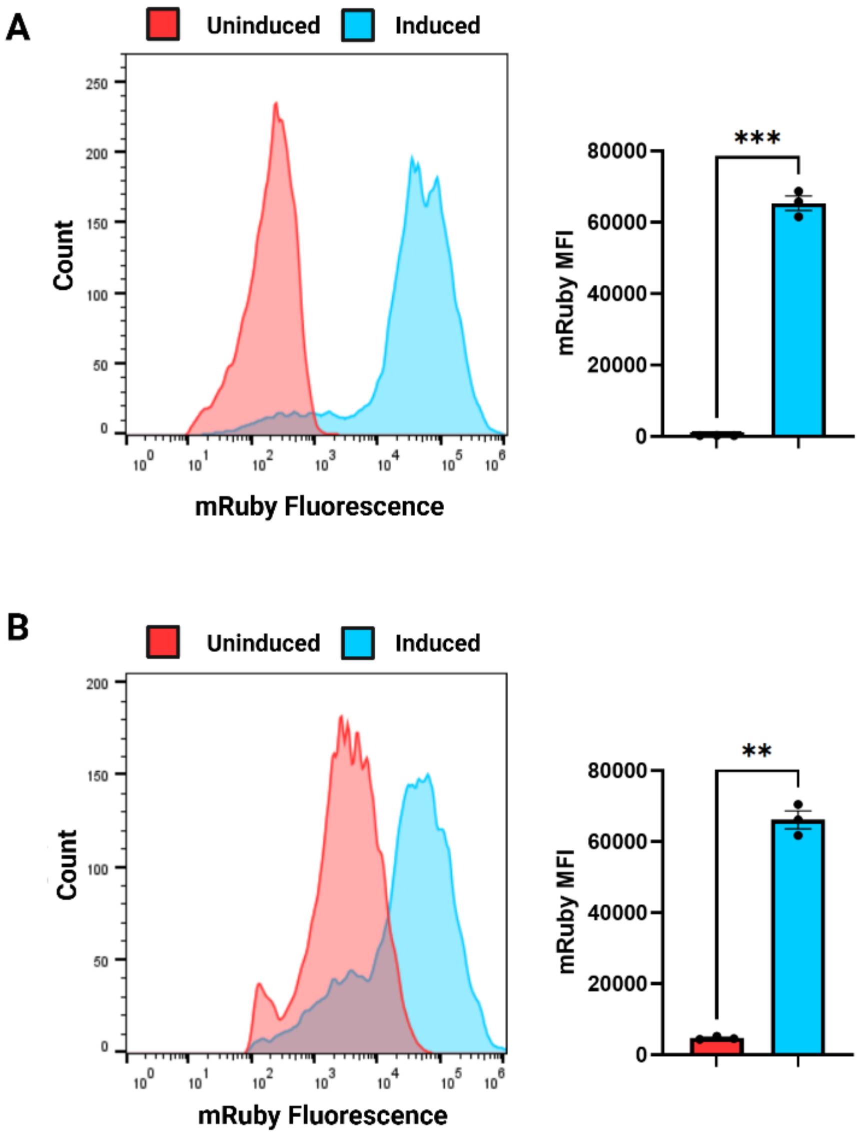


**Supplemental Figure 2: Testing IPTG and Tetracycline Transcription Factors in BIT Yeast Strain**. **A)** A plasmid with mRuby fluorescent protein DNA under a pLacFec promoter was transformed into the BIT yeast strain and uninduced and induced populations were compared. **B)** A plasmid with mRuby fluorescent protein DNA under a pTet7 promoter was transformed into the BIT yeast strain and uninduced and induced populations were compared. Flow cytometry data comparing the uninduced and induced populations of this construct are included (n=3). *P* values were determined by using a paired t-test between the uninduced and induced samples. All data are shown as mean±s.e.m
